# *Staphylococcal* sRNA IsrR down-regulates methylthiotransferase MiaB under iron-deficient conditions

**DOI:** 10.1101/2023.11.09.566390

**Authors:** Maxime Barrault, Elise Leclair, Etornam Kofi Kumeko, Eric Jacquet, Philippe Bouloc

## Abstract

*Staphylococcus aureus* is a major contributor to bacterial-associated mortality, owing to its exceptional adaptability across diverse environments. Iron is vital to most organisms but can be toxic in excess. To manage its intracellular iron, *S. aureus*, like many pathogens, employs intricate systems. We have recently identified IsrR as a key regulatory RNA induced during iron starvation. Its role is to reduce the synthesis of non-essential iron-containing proteins under iron-depleted conditions. In this study, we unveil IsrR’s regulatory action on MiaB, an enzyme responsible for methylthio group addition to specific sites on transfer RNAs (tRNAs). We use predictive tools and reporter fusion assays to demonstrate IsrR’s binding to the Shine-Dalgarno sequence of *miaB* RNA, thereby impeding its translation. The effectiveness of IsrR hinges on the integrity of a specific C-rich region. As MiaB is non-essential and has iron-sulfur clusters, IsrR induction spares iron by downregulating *miaB*. This may improve *S. aureus* fitness and aid in navigating the host’s nutritional immune defenses.

**IMPORTANCE:** In many biotopes, including those found within an infected host, bacteria confront the challenge of iron deficiency. They employ various strategies to adapt to this scarcity of nutrients, one of which involves regulating iron-containing proteins through the action of small regulatory RNAs. Our study shows how IsrR, a small RNA from *S. aureus*, prevents the production of MiaB, a tRNA-modifying enzyme containing iron-sulfur clusters. With this illustration, we propose a new substrate for an iron-sparing small RNA, which, when downregulated should reduce the need for iron and save it to essential functions.

## INTRODUCTION

*Staphylococcus aureus* is the leading cause of bacteria-associated mortality worldwide (1). Its pathogenicity is due to its adaptability across various biotopes and the expression of multiple virulence determinants. Iron is a trace element universally required for growth, but toxic in excess. *S. aureus*, akin to other pathogens, has elaborate systems for maintaining its intracellular iron homeostasis to match environmental conditions (2).

We recently identified and characterized IsrR, a small regulatory RNA (sRNA) of *S. aureus*, contributing to an adaptive response to iron scarcity (3, 4). Most sRNAs exert their activity by pairing with target mRNAs whose stability and/or translation they affect. They are generally expressed conditionally and contribute to adaptation to various growth conditions. Their estimated number in *S. aureus* varies from around 50 to a few hundred according to studies. They are major regulators or contributors to fine-tune processes including metabolism, virulence, and antibiotic resistance; however, the function of most of them remains unknown (5–7).

IsrR, by restricting the production of non-essential iron-containing proteins, is expected to spare iron for indispensable functions. *isrR* expression is repressed by the ferric uptake regulator (Fur) which is active in iron-rich environments. Induced during iron deficiency, IsrR targets Shine-Dalgarno motifs on mRNAs coding for iron-containing proteins, notably restraining translation of specific mRNAs (*fdhA*, *narG*, *nasD*, and *gltB2*) linked to nitrate respiration and encoding [Fe-S] cluster-dependent enzymes (3). IsrR also affects the TCA cycle by down-regulating aconitase and its positive regulator CcpE (4). Given limited iron requirements, fermentative pathways are favored under iron-limited conditions. IsrR is a functional analog of RyhB from *Escherichia coli* (8) and of other sRNAs found in diverse bacterial species (reviewed in (9, 10)) sharing targets and iron regulation. However, IsrR does not share sequence similarities or the same enzymatic requirements for its activity with RyhB and its reported functional analogs. IsrR, conserved within the Staphylococcus genus, is likely not an ortholog of these sRNAs, pointing to a convergent evolutionary solution for adaptation to low-iron conditions (3).

Hosts prevent pathogen growth through an arsenal of defense mechanisms, one of which, known as nutritional immunity, is the sequestration of nutrients such as iron (11). We reported that IsrR deficiency results in reduced virulence of *S. aureus* in a murine infection model, highlighting the role of IsrR in facilitating optimal spread of *S. aureus* in the host environment, probably by facilitating evasion of nutritional immunity (3).

Here, we demonstrate that IsrR downregulates *miaB* encoding a non-essential protein containing [Fe-S] clusters. This new example further supports the view that IsrR is a central player in the maintenance of iron homeostasis in *S. aureus*.

## MATERIALS AND METHODS

### Bacterial strains, plasmids and growth conditions

Bacterial strains, plasmids and primers used for this study are described in Table 1. Plasmids were engineered by Gibson assembly (12) in *E. coli* IM08B (13) as described (Table 1), using the indicated appropriate primers for PCR amplification. Plasmids were verified by DNA sequencing and then transferred into *S. aureus*.

**Table 1.**
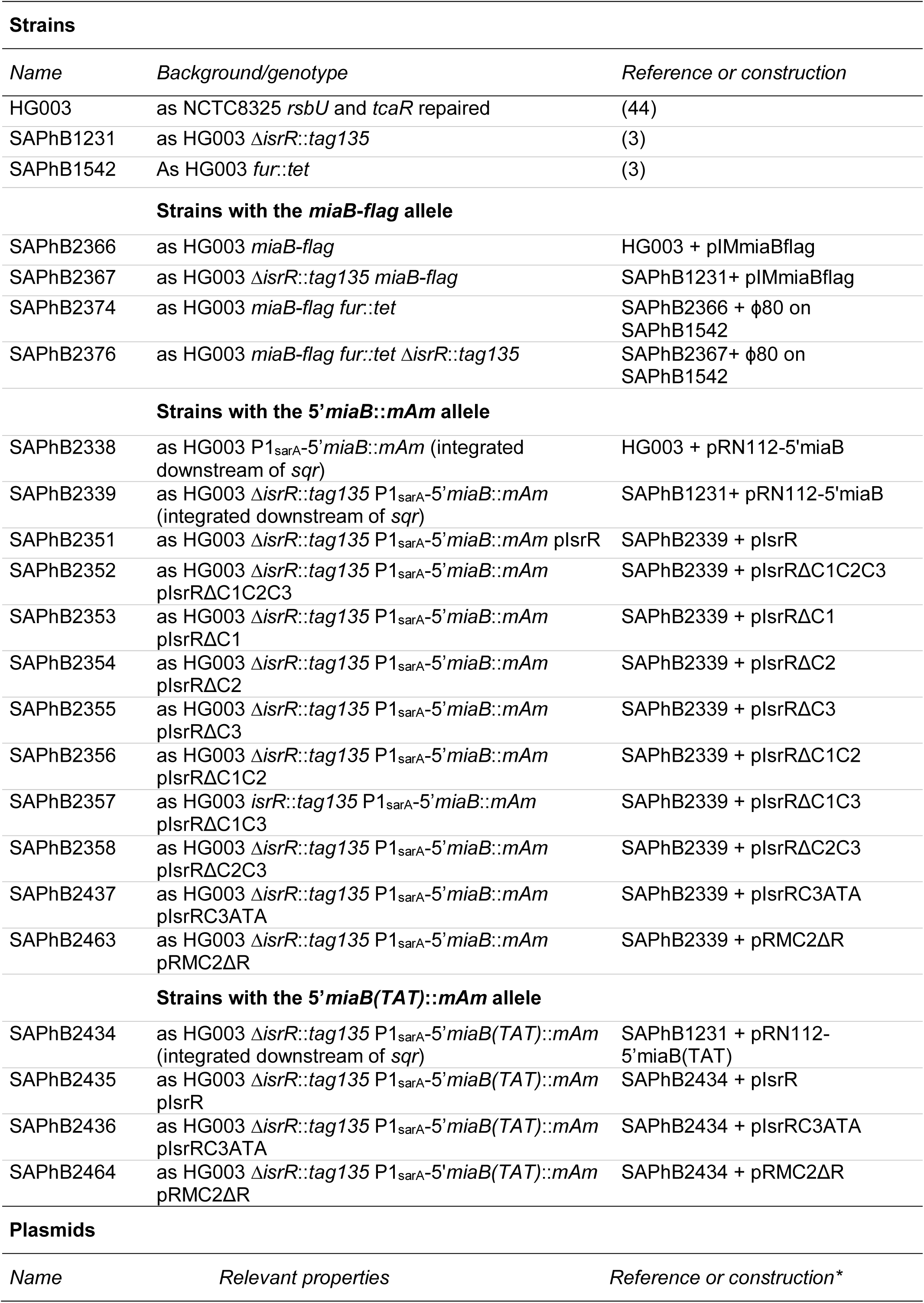
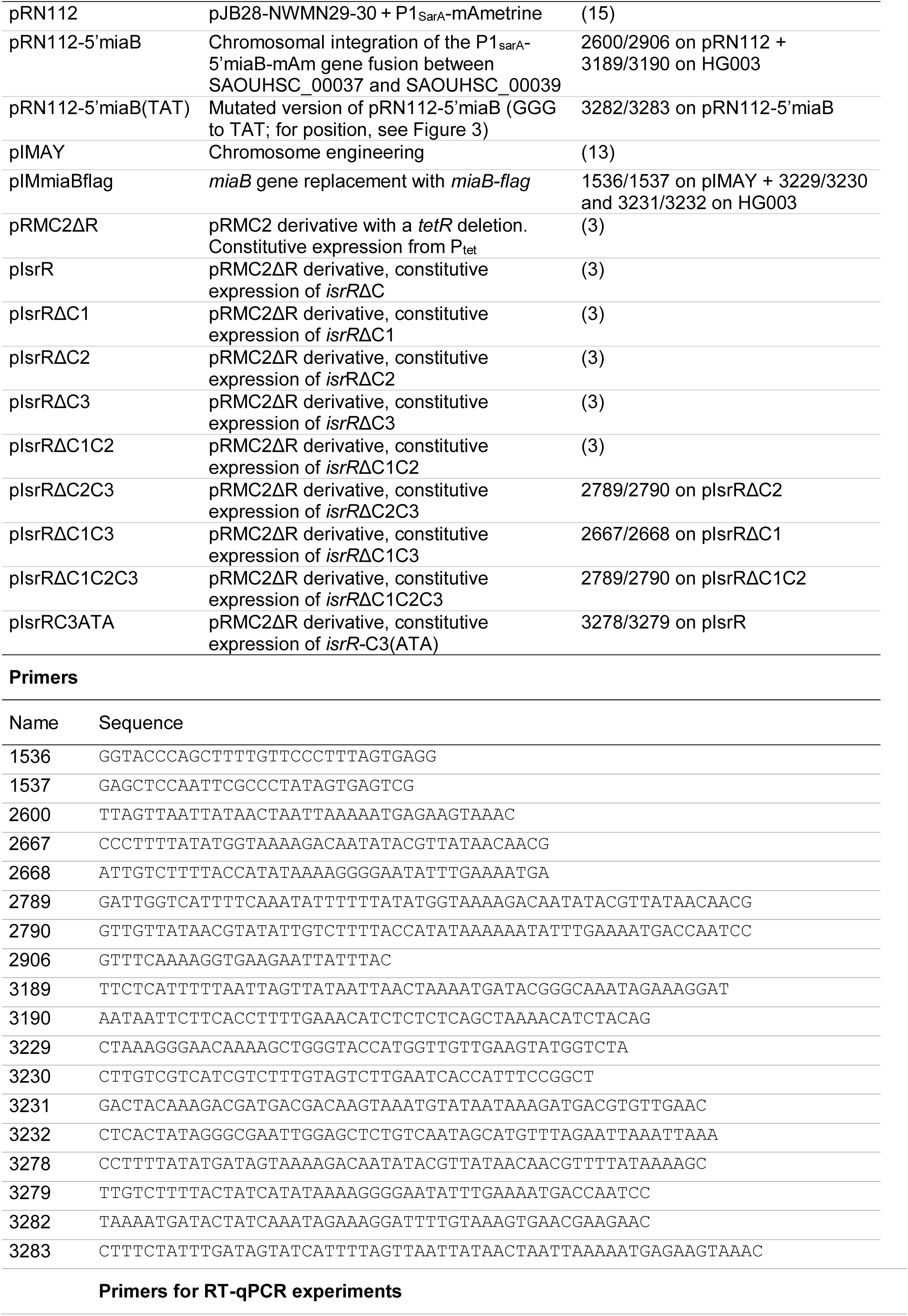
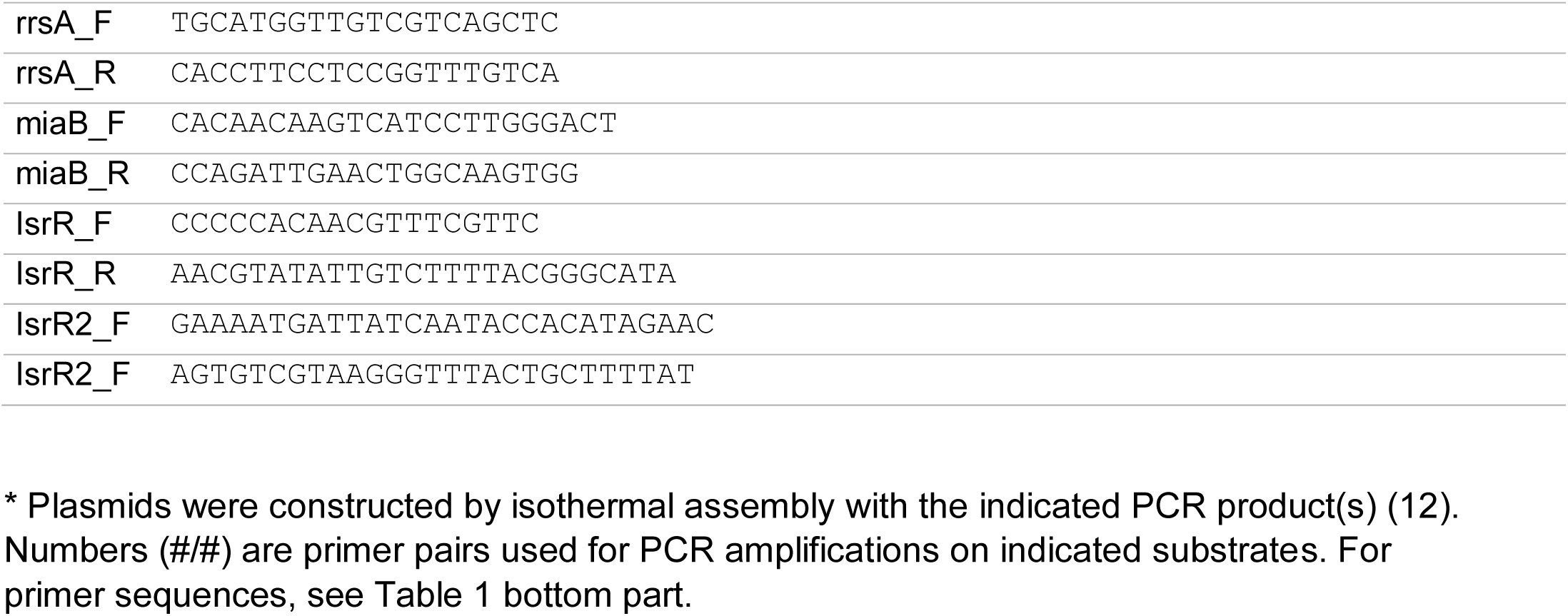
Strains, plasmids, and DNA primers used in this study.

Strains in which the chromosomal *miaB* gene was replaced by the *miaB-flag* allele were constructed using pIMmiaBflag, a pIMAY (13) derivative, as described (14). Strains carrying the chromosomal reporter gene 5’*miaB-mAm* and its mutated derivative 5’*miaB(TAT)-mAm* inserted downstream *sqr* were constructed using the pRN112 derivatives pRN112-5’miaB and pRN112-5’miaB(TAT), respectively, as described (15). The *fur*::*tet* allele was transferred by ϕ80 phage-mediated transduction.

*S. aureus* strains were grown aerobically in Brain Heart Infusion (BHI) broth at 37°C. *E. coli* strains were grown aerobically in lysogenic broth (LB) at 37°C. Antibiotics were added to media as needed: ampicillin 100 μg/ml and chloramphenicol 20 μg/ml for *E. coli*; chloramphenicol 5 μg/ml and tetracycline 2.5 μg/ml for *S. aureus*. Iron-depleted media was obtained by the addition of DIP (2,2’-dipyridyl) 0.5 mM incubated for 30 min before any use.

### Biocomputing analysis

Pairing predictions between IsrR and the mRNA targets were made using IntaRNA (16) set with default parameters except for suboptimal interaction overlap set to “can overlap in both”. The sequences used for IsrR and mRNA targets were extracted from the *S. aureus* NCTC8325 strain (NCBI RefSeq NC_007795.1). For the mRNA targets, the sequences used started at the TSS when known (e.g., Exact Mapping Of Transcriptome Ends, EMOTE (17) or were arbitrarily made to start at nucleotide -60 with respect to the +1 of translation. The sequences contain the 5’ UTR and the 17 first codons of the CDS.

### Statistical tests

For Western-blots, statistical analyses between two groups were made by testing the equality of variances between the groups using a Fisher test and, if and only if the two variances are equal, comparing them using a Student t-test. For fluorescence experiments, significant differences in the means between different groups were assessed by an ANOVA (or Welch ANOVA in case of inequality of variances) to identify significantly different groups. Error bars represent the standard deviation of the results.

### Western blot analysis

The chromosomal *miaB-flag* reporter gene was constructed as described (Table 1). Strains were grown overnight with and without DIP. Protein extracts were prepared as follows: 4 mL of the bacterial culture were harvested by centrifugation at 4°C and 3800 g for 5 min. The resulting pellet was resuspended in 400 µL of 50 mM Tris-HCl. Cell disruption was achieved using a Fast-Prep homogenizer (MP Biomedicals). After disruption, crude protein extracts were obtained by centrifugation at 4°C and 21000 g for 15 min. The protein concentration of each extract was determined using the Bradford protein assay method. Ten µg of each protein extract were loaded onto a NuPAGE 4%-12% Bis-Tris gel (Invitrogen). Electrophoresis was carried out at 150V for 45 min. Proteins were transferred from the gel to a PVDF membrane using the iBind system (Invitrogen). Immunolabeling was performed overnight using a rabbit anti-Flag antibody (Invitrogen) as primary antibody and an anti-rabbit IgG HRP-conjugated antibody (Promega) as secondary antibody. The iBlot system (Invitrogen) was employed for antibody incubation. Images of the immunolabeled membrane were acquired using a ChemiDoc MP imaging system (Bio-Rad). Bands were quantified using the Image Lab software (Bio-Rad).

### Fluorescent translational reporter assay

A *miaB* translational reporter (P_sarA_-5’miaB-mAm) was inserted between genes *SAOUHSC_00037* and *SAOUHSC_00039* in HG003 and HG003 Δ*isrR*::*tag135* using pRN112-5’miaB (Table 1), a pRN112 derivative, as described (15). Notably, the mAmetrine reporter gene (*mAm*) was placed in-frame and downstream *miaB* 18^th^ codon with the *miaB* 5’UTR placed under the control of the P1 promoter of *sarA* (P1_sarA_). Subsequently, the strains carrying the reporter fusion were transformed with plasmids expressing *isrR*, along with its various derivatives featuring CRR deletions. These plasmids, pIsrR, pIsrRΔC1, pIsrRΔC2, pIsrRΔC3, pIsrRΔC1C2, pIsrRΔC1C3, pIsrRΔC2C3, and pIsrRΔC1C2C3, are pRMC2ΔR derivatives (3, 4). For fluorescence measurements, the transformed strains were cultured overnight in BHI medium supplemented with chloramphenicol and DIP. Subsequently, cultures were washed three times with 1x phosphate-buffered saline (PBS). Fluorescence levels were quantified using a microplate reader (CLARIOstar) with excitation at 425 nm and emission at 525 nm. To ensure accuracy, fluorescence measurements were normalized by subtracting the auto-fluorescence of the HG003 strain (lacking the reporter construct), and all cultures were adjusted to an optical density at 600 nm (OD_600_) of 1.

### Quantitative reverse transcriptase PCR

To determine the stability of *miaB* mRNA, overnight cultures HG003 and HG003 Δ*isrR* (biological triplicates, N=3) were diluted to an OD_600_ = 0.005 and incubated in BHI supplemented with DIP (0.5 mM) at 37°C. At OD_600_ = 1, rifampicin (concentration 200 µg/ml) was added to the growth medium. Bacterial cultures were harvested before rifampicin addition (time 0) and at 2, 5, and 10 min after rifampicin addition. Total RNAs were extracted as described (18) and treated with DNase (Qiagen) according to manufacturer’s instructions. The last purification step was performed with the RNA Nucleospin kit according to manufacturer’s instructions (Macherey-Nagel). The integrity of the RNAs was verified using the Agilent 2100 bioanalyzer with the RNA 6000 Nano kit (Agilent Technologies). RT-pPCR experiments and data analysis were performed as described (19). The Ct values of *rrsA* were used to normalize the data and the determination of the gene expression ratio was achieved using the ΔΔCt method. Primers used for RT-qPCRs are indicated in Table 1.

IsrR and its variant expressed from plasmids were quantified as indicated above except that total RNAs were extracted from overnight cultures. Since isrR derivatives have different mutations, a different set of primers was used (Table 1, primer section; IsrR2_F and IsrR2-R).

## RESULTS

### MiaB is a putative IsrR-target

We used CopraRNA (20), a software designed for predicting targets of bacterial sRNAs across diverse species, to unveil potential targets of IsrR. Given *isrR* conservation throughout the Staphylococcus genus, CopraRNA emerged as an efficient tool for IsrR target prediction (21), resulting in the successful validation of six of its targets as reported (3, 4). Among the predicted targets was *miaB* mRNA (3, 22). The latter encodes MiaB, a methylthiotransferase catalyzing the addition of a methylthio group (ms^2^) to isopentenyl adenine (i^6^A) at position 37 (ms^2^i^6^A-37) of transfer RNA (tRNA) anticodons (23). This modification is present in most tRNAs reading codons starting with a U and is conserved from bacteria to human (24); it contributes to stabilizing the codon–anticodon interaction (25) and affects translational accuracy (26).

To substantiate the proposed interaction between IsrR and *miaB* mRNA, we used the IntaRNA software (16). The input was the complete IsrR sequence and the *miaB* 5’ untranslated region (UTR) with 25 codons. IsrR is characterized by three single-stranded C-rich regions, denoted as CRR1, CCR2, and CRR3 (3). The CRRs are common sequences among Firmicutes’ regulatory RNAs where they are presumably seed-binding motifs to G-rich sequences (27). IntaRNA analysis suggests a potential pairing of IsrR with the Shine-Dalgarno sequence of *miaB* RNA (Figure 1A). The pairing involves 55 nucleotides, an unusually high number of pairs. The energy of sRNA-mRNA pairing is low (-22.33 kcal/mol) supporting its existence. It involves the CCR3 GCCCG sequence, which could serve as a seed motif of the interaction. Of note, the isrR / *miaB* mRNA pairing also is conserved within the Staphylococci genus (Figure 1B).

**Figure 1:**
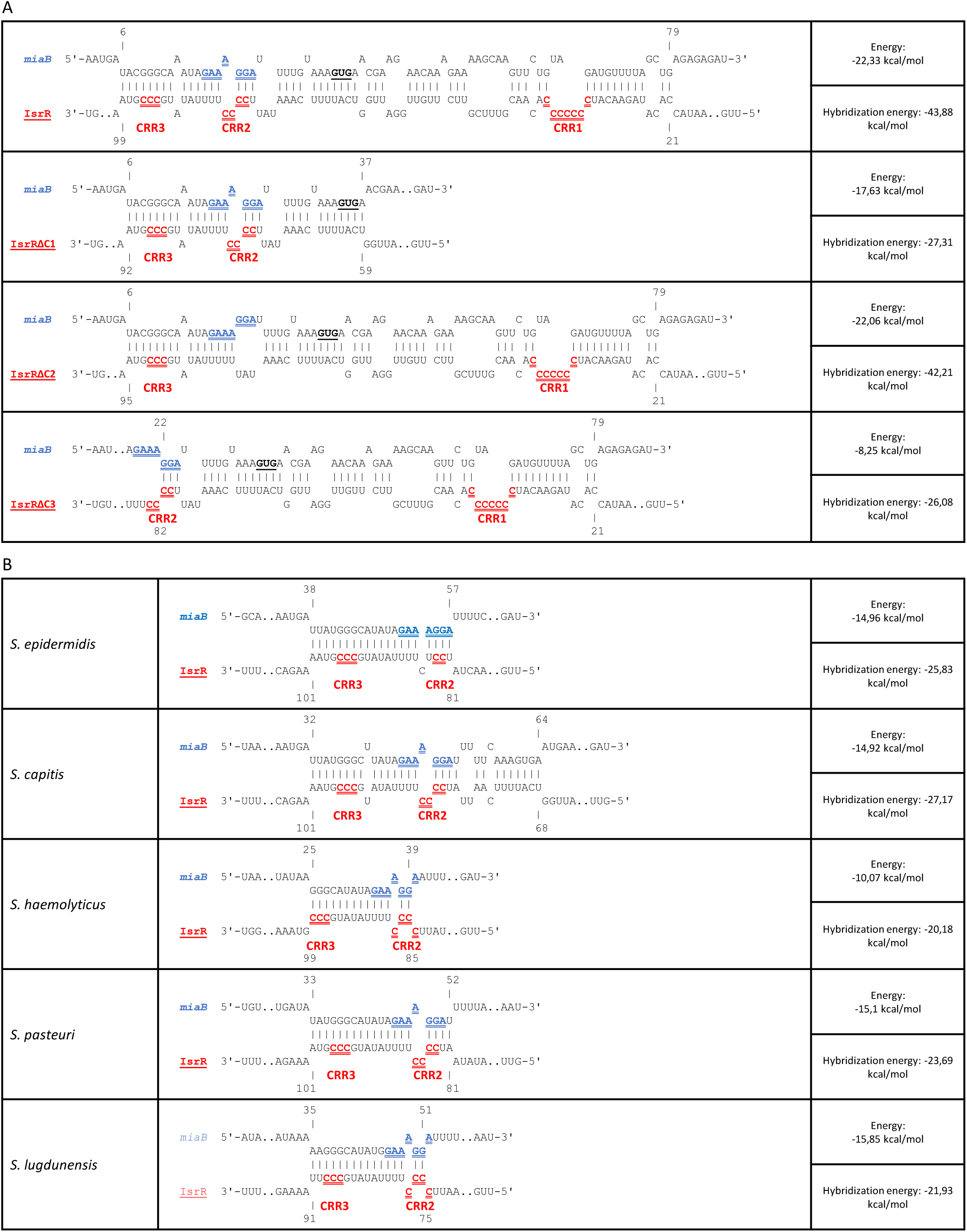
Computer prediction of IsrR and mutated IsrR pairing with *miaB* mRNA. A) IntaRNA (16) pairing prediction of IsrR and IsrR derivatives deleted for CRR1, CRR2 or CRR3 with *miaB* mRNA. Blue sequences, SD; red sequences, CRRs; bold underline characters, GUG start codon; E, energy; HE, hybridization energy. B) IntaRNA pairing prediction of IsrR and *miaB* mRNA in different species from the *Staphylococcus* genus. IsrR sequences from the indicated species were obtained with GLASSgo (45). For legends, see Figure 1A.

### IsrR-dependent reduction of MiaB expression

To explore the potential influence of IsrR on *miaB* expression, a gene fusion was constructed wherein the native *miaB* gene was replaced by a copy harboring the complete *miaB* open reading frames extended with a C-terminal Flag sequence (Figure 2A). The resulting MiaB-Flag fusion protein was detectable through anti-Flag antibodies, thereby serving as a quantifiable surrogate for wild-type gene expression levels. This reporter fusion was integrated into the chromosomes of both HG003 and its corresponding Δ*isrR* isogenic derivative. Western blot analyses conducted on HG003 cultures cultivated under nutrient-rich conditions show an unaltered MiaB-Flag abundance, irrespective of the presence or absence of *isrR*. This status quo was in line with the known Fur-mediated repression of *isrR* under iron-replete conditions. We previously reported that introducing the iron chelator 2,2’-dipyridyl (DIP) into the growth medium elicits a release from Fur repression, consequently inducing IsrR expression (3). Remarkably, under DIP-induced conditions, no MiaB-Flag was detected regardless of the *isrR* background (Figure 2B). As iron depletion affects [Fe-S] clusters in proteins, we hypothesized that the presence of DIP could lead to the formation of apo-MiaB possibly unstable.

**Figure 2:**
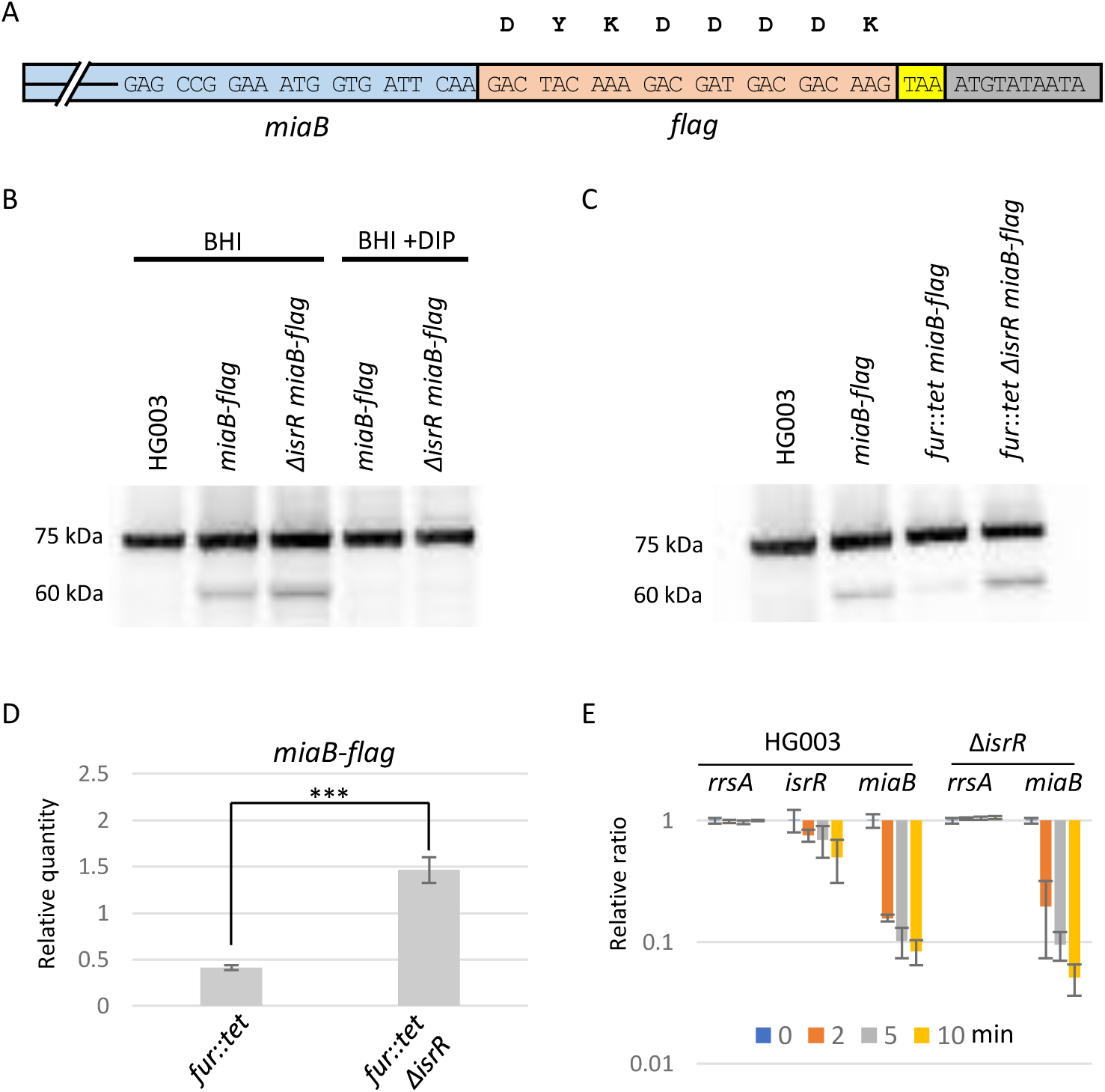
MiaB-Flag reporter detection. A) Schematic representation of the MiaB-Flag reporter fusion. Blue, 3’ end of *miaB* ORF; Orange, *flag* sequence; Yellow, native stop codon; Grey, first nts of *miaB* 3’ UTR. B) Western blot experiment with anti-Flag antibodies in presence or absence of iron chelators (N=4). Genotypes and growth conditions are indicated. 75 kDa, non-specific signal present in all samples including Flag-less strain. 60 kDa, signals corresponding to MiaB-Flag. C) Western blot experiment with anti-Flag antibodies with *fur* and *fur*::*tet* strains (N=3) as explained in Figure 2B. D) Histograms. MiaB-Flag signals from Western blot (Figure 2C) were normalized to the 75 kDa signals from their corresponding lane. Histogram y-axis indicates the values of normalized MiaB-Flag signals of indicated strains divided by the normalized MiaB-Flag signal from the wild-type background strain (*fur*^+^). Error bars indicate the standard deviation of three biological replicates. ***, P-value < 0,001 for Student test (N=3). E) The stability of *miaB* mRNA is not significantly affected by IsrR. HG003 and its isogenic Δ*isrR* derivative were grown in rich medium supplemented with DIP. At t0, rifampicin (200 µg/ml) was added to the growth medium. Cultures were sampled at t0, 2, 5, and 10 min after the addition of rifampicin, total RNA was extracted, and the amounts of *miaB* mRNA, *rrsA* rRNA (control), and IsrR were determined by RT-qPCR. Histograms Y-axis show the quantification of *miaB* mRNA, *rrsA* sRNA, and IsrR normalized to t0. Error bars indicate the standard deviation of biological triplicates (N=3).

To overcome this issue, a *fur*::tet allele (28) was introduced by phage-mediated transduction in the *miaB-flag* and Δ*isrR miaB-flag* strains rendering *isrR* constitutively expressed, as previously reported (3). Expectedly, in this genetic background, MiaB-Flag was barely detected in the *fur* strain as opposed to its *fur* Δ*isrR* derivative (Figure 2C-D).

### IsrR-mediated post-transcriptional regulation of *miaB* expression

In the absence of Fur, *isrR* is expressed and *miaB* is downregulated; the observation is explained by predicted base pairing interaction between IsrR and *miaB* RNA. In many cases, sRNAs targeting mRNAs affect their stability. We therefore considered that IsrR might decrease *miaB* mRNA stability. This question was addressed by comparing *miaB* mRNA stability in wild-type and Δ*isrR* strains under iron-starved condition. At OD_600_=1, RNA synthesis was inhibited by rifampicin addition to the growth medium, and *miaB* mRNA turnover was determined by RT-qPCR at different time points. No significant differences were observed, suggesting that IsrR does not affect *miaB* mRNA stability (Figure 2E). We, therefore, considered that the regulation of *miaB* by IsrR occurred primarily at the translational level, a feature already observed for the regulation of *fdhA* and *gltB2* by IsrR (3). To examine this hypothesis, a genetic reporter of *miaB* translation was constructed. It is based on pRN112, a replication thermosensitive plasmid for chromosomal integration carrying the mAmetrine gene (*mAm*) under the transcriptional control of the P1 *sarA* promoter (P1_sarA_) (15). The 5’ untranslated region (5’UTR) of *miaB*, along with an additional 54 nucleotides, was inserted within pRN112 under the transcriptional control of P1_sarA_. The first 18 *miaB* codons were positioned upstream and in frame *mAm* (Figure 3A). With this construct, the mAmetrine fluorescence served as a proxy for *miaB* expression levels, and as the reporter gene is under P1_sarA_ transcriptional regulation. The transcription from this promoter is expected to be constitutive therefore not associated with any IsrR-dependent transcriptional regulation. To prevent concerns associated with multi-copy reporter plasmids, the engineered sequence was integrated into the chromosome of both HG003 strain and its Δ*isrR* derivative. As for the endogenous *miaB*, the 5’*miaB*-mAm reporter exhibited significant downregulation under iron-starved conditions in an IsrR-dependent manner, while its transcription was driven from P1_sarA_ (Figure 3B).

**Figure 3:**
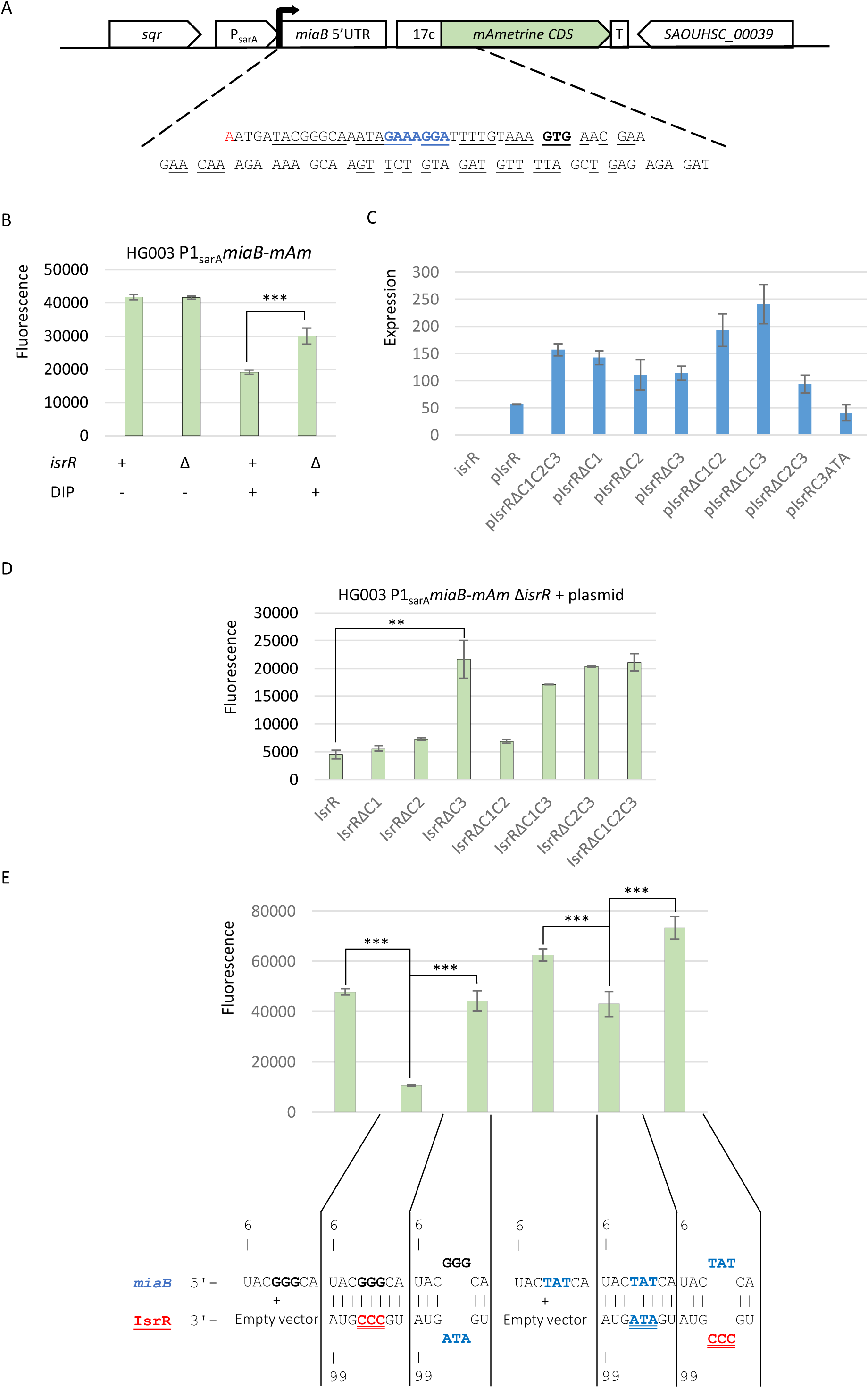
Translational repression of *miaB* reporter fusion by IsrR. A) Chromosomal reporter fusion for detection of IsrR activity. Red nt; transcription start site, blue bold nts; RBS, black bold nts; start codon, underlined nts; nts in interaction with IsrR as predicted by IntaRNA. B) Translational activity of the *miaB-mAm* reporter fusion (mAmetrine fluorescence signal) in HG003 and its Δ*isrR* derivative grown in BHI and BHI supplemented with DIP as shown. Results (N=3) are normalized to OD_600_=1. C) Expression of *isrR* and its derived alleles expressed from the indicated plasmids in stationary phase normalized to wild-type *isrR* expression (first histogram). The amount was determined by RT-qPCR. Error bars indicate the standard deviation of four independent biological replicates, with the exception of pIsrRΔC3, which has three. D) The Δ*isrR* strain harboring the *miaB-mAm* reporter fusion was transformed with plasmids expressing different versions of IsrR and cultured in BHI supplemented with DIP. The translational activity of the reporter fusion was quantified by measuring the fluorescence. The significance of the difference in fluorescence between IsrR and IsrRΔC3 groups is supported by ANOVA: results (N=3) are normalized to OD_600_=1. **, P-value < 0.01; ***, P-value < 0.001. E) The Δ*isrR* strains harboring the *miaB-mAm* reporter fusion or its mutated derivative in its three G motif (GGG to TAT) were transformed with an empty plasmid, a plasmid expressing IsrR or a plasmid expressing *isrR* mutated in CRR3 (CCC change to ATA). Mutations in the three G motif and CCR3 were designed to be complementary as shown below the histograms. Strains were cultured in BHI supplemented with DIP. The translational activity of the reporter fusions, represented by histograms, was quantified by measuring the fluorescence (N=5). The significance of the difference in fluorescence between the optimum pairing and altered pairing is supported by ANOVA and Welch tests.

Our experimental findings support that IsrR downregulates the expression of the reporter fusion by a post-transcriptional event.

### The C-rich region 3 (CRR3) of IsrR plays a pivotal role in the downregulation of *miaB* **mRNA**

Pairing interactions between sRNAs and their targets are initiated by a kissing complex between single-stranded regions, which then propagates within both partners (29). sRNAs can have multiple seed regions, usually C-rich in *S. aureus* (27), specific to given targets as exemplified by RNAIII (review in (5)) or with redundant activities as in RsaE (30). To determine specific regions of IsrR responsible for its regulatory activity against *miaB* mRNA, experiments using the HG003 Δ*isrR* strain, harboring the 5’*miaB* fluorescent reporter were conducted. Plasmids expressing various IsrR derivatives, including wild-type IsrR, IsrR lacking CRR1, CRR2, CRR3, or all three CRRs were used. *isrR* was placed under the control of the *tet* promoter (P_tet_) in the absence of the Tet repressor (TetR), although it should be noted that a Fur binding site is present within the transcribed sequence (3). Consequently, we supplemented the growth media with DIP to prevent Fur repression on *isrR* expression. All plasmids led to overexpression of IsrR and derived alleles (compared to the expression from the wild-type *isrR* endogenous copy) under the tested condition, stationary phase culture in rich medium supplemented with DIP (Figure 3C). Their quantity varies from one allele to another, probably due to different stability, but remains largely sufficient to be potential active.

The presence of a plasmid expressing IsrR markedly diminished the fluorescence of the reporter fusion when compared to a control plasmid or a plasmid expressing IsrR with no CRR motif. In contrast, the deletion of CRR3 resulted in a significant increase in mAmetrine fluorescence indicating a loss of activity against the reporter fusion. Note that the same construct producing IsrRΔC3 is expressed and active against *gltB2* and *fdhA* mRNAs (3). However, the deletion of CRR1 and CRR2 in isolation had no substantial impact on IsrR’s activity against the 5’*miaB* reporter (Figure 3C). This observation suggests that these two motifs may either be dispensable for the regulatory activity or possess redundant functions with respect to the *miaB* 5’UTR.

To discern between these possibilities, we introduced into the Δ*isrR*::*tag135* strain plasmids expressing different *isrR* derivatives with deletions of two CRRs. As expected, strains carrying alleles with the CRR3 deletion (*isrR*ΔC1C3 and *isrR*ΔC2C3) failed to complement the Δ*isrR* allele. In contrast, *isrR*ΔC1C2 was still capable of downregulating the fluorescence of the P_sarA_*5’miaB-mAm* reporter (Figure 3C). These findings align with the results obtained from IntaRNA analysis, which indicates that the *IsrR*/*miaB* mRNA pairing energy remains substantial as long as CRR3 and its surrounding region remain intact (Figure 1). To further support the involvement of CRR3 in *IsrR*/*miaB* mRNA pairing, the reporter system was tested with point mutations altering the predicted pairing. First, the CRR3 motif was changed from CCC to ATA leading to IsrR-C3(ATA). Expectedly, mutated IsrR prevented the *miaB* reporter fusion downregulation (Figure 3E). A compensatory mutation restoring the pairing with the mutated IsrR was introduced within the *miaB* sequence by changing a GGG sequence upstream *miaB* Shine Dalgarno sequence to TAT. These mutations altering the 5’UTR region led to a slightly higher level of reporter gene expression. However, while wild-type IsrR did not affect the expression of the mutated reporter fusion, mutated-IsrR (IsrR-C3(ATA)) downregulated its expression. Down-regulation of miaB(TAT) by IsrR-C3(ATA) was less marked than with *isrR* and *miaB* wild-type sequences, perhaps due to an altered pairing structure or the fact that *isrR-C3*(*ATA*) is less expressed than *isrR* (Figure 3C). These experiments support the conclusion that CRR3/5’UTR *miaB* RNA pairing is needed for the IsrR-dependent downregulation of *miaB*.

It is worth noting that the requirement for CRRs in IsrR activity appears to be contingent upon its target mRNA (Figure 4). For instance, the absence of CRR3 does not impede the downregulation of *gltB2* and *fdhA* mRNAs (3). Conversely, for the latter mRNAs, the integrity of both CRR1 and CRR2 is indispensable, presenting a contrast to *miaB* mRNA regulation. This observation suggests that the acquisition of multiple CRRs, together with their adjacent regions, confers on regulatory RNAs the ability to control a wider range of target mRNAs. sRNAs likely evolve by acquiring new domains to interact with different targets (*e.g. miaB* RNA vs *gltB2* mRNA). At the same time, mRNAs may evolve to adapt to the available sRNA motifs (*e.g. gltB2* RNA vs *fdhA* mRNA).

**Figure 4:**
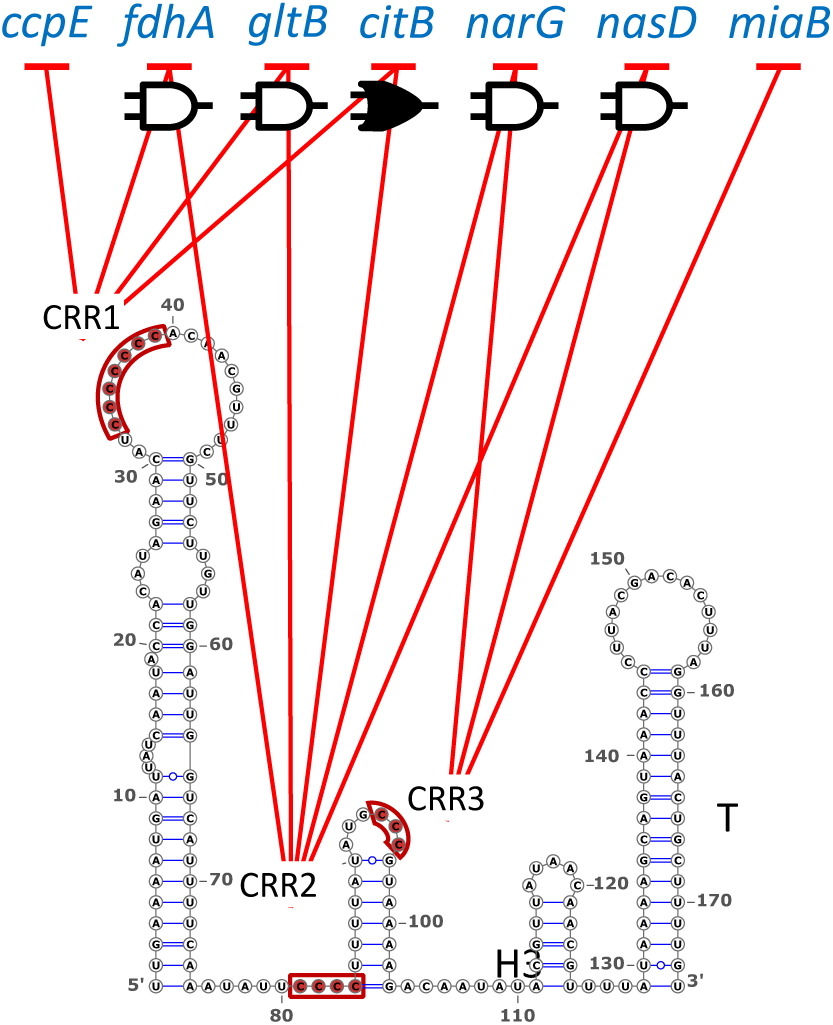
Contribution of IsrR CRRs to downregulate different mRNA targets. Negative arrows starting with CRR1, 2 or 3 and pointing to mRNA targets indicate CRRs required for IsrR activity towards the corresponding targets. Boolean symbols indicate that regulation requires the integrity of two CRRs (AND gate, empty symbol) or that two CRRs can act independently of each other (OR gate, solid symbol). Data derived from results obtained with fluorescent reporters in (3, 4) and this work.

## DISCUSSION

MiaB is a conserved enzyme that modifies tRNAs acting after MiaA on an adenine adjacent to the anticodon site (23–26). In some species, the modified adenosine is then hydroxylated by MiaE, but this enzyme is not present in *S. aureus* (31, 32). RNA modifications optimize codon/anticodon interactions and improve translation fidelity; for example, the absence of MiaB was shown to affect the efficiency of a suppressor tRNA (23).

The relationship between iron metabolism and the MiaB enzyme has been a subject of scientific interest for several decades. As far back as the 1960s, researchers noted the absence of certain tRNA modifications in *E. coli* cultured in iron-free media, underscoring the connection between iron availability and tRNA modifications (33). Subsequent investigations revealed that the *miaB* gene encodes the tRNA methylthiotransferase responsible for these modifications, and mutations in *miaB* were found to reduce decoding efficiencies under specific conditions (23). Notably, MiaB was initially thought to contain two [4Fe-4S] clusters (34), but a more recent study concludes that MiaB has one [4Fe-4S] and one [3Fe-4S] (35). Under iron-starved conditions, clusters depletion renders MiaB inactive. Furthermore, as shown here, in iron-starved condition, MiaB-Flag was not detectable suggesting that in the absence of iron cluster, MiaB could be degraded as observed for other proteins with [Fe-S] clusters (36). For example, the glutamine phosphoribosylpyrophosphate amidotransferase of *Bacillus subtilis* has a [4Fe-4S] cluster which when altered by O_2_ favors enzyme degradation (37).

In *E. coli*, the influence of MiaB extends to the regulation of Fur through a distinctive mechanism (38). The *fur* gene is preceded by *uof*, an open reading frame located “<u>u</u>pstream <u>o</u>f fur”, and the translation of *fur* is tightly coupled to that of *uof*. An intriguing aspect of this regulatory relationship is the presence of a codon UCA, decoded by a MiaB-dependent tRNA^Ser^. Consequently, efficient *uof*/*fur* translation relies on functional MiaB. When iron-starvation conditions prevail, MiaB becomes inactive, impairing tRNA methylthiolation. This, in turn, leads to reduced *uof*/*fur* translation, ultimately favoring the expression of iron uptake genes. However, this regulation is not conserved in *S. aureus* since no *uof*-like sequence was observed.

In standard laboratory conditions, Enterobacteria appear to be resilient to the absence of MiaB (39). In the Gram-positive bacterium *B. subtilis*, *miaB* (aka *ymcB*) is also non-essential under the tested conditions (40) while its inactivation is associated with the disappearance of ms^2^i^6^A post-transcriptional modifications (41). There is a 68% amino acid identity between *B. subtilis and S. aureus* MiaBs, suggesting that Staphylococcal MiaB indeed methylthiolates tRNAs. *miaB* in *Staphylococcus aureus* is also not essential (42). Consequently, preventing MiaB activity is likely to have minimal consequences in various growth conditions. Since MiaB needs [Fe-S] clusters as cofactors, suppressing its synthesis in iron-starved environments spares iron, allowing it to be redirected toward essential cellular processes.

It is noteworthy that the inhibition of *miaB* RNA translation by a sRNA associated with the iron-sparing response is a feature observed thus far only in *S. aureus*. Surprisingly, in *E. coli*, RyhB does not target *miaB* mRNA (43), presumably because MiaB function is more critical to some cellular functions in this organism. The absence of MiaB can result in reduced translation accuracy, impacting gene expression. Therefore, modulating *miaB* expression to spare iron represents a viable alternative for bacteria, provided the fitness benefits outweigh the costs.

## ACKNOWLEDGMENTS

We thank our lab colleagues and Max Ipane for helpful discussions. This work was funded by the *Agence Nationale pour la Recherche* (ANR) ANR grant [ANR-19-CE12-0006-01 (sRNA-RRARE)] and by the *OI Microbes* from the *Université Paris-Saclay*. MB was the recipient of fellowships from the *ministère de l’Enseignement Supérieur et de la Recherche*.

